# Spatial interactions modulate tumor growth and immune infiltration

**DOI:** 10.1101/2024.01.10.575036

**Authors:** Sadegh Marzban, Sonal Srivastava, Sharon Kartika, Rafael Bravo, Rachel Safriel, Aidan Zarski, Alexander Anderson, Christine H. Chung, Antonio L. Amelio, Jeffrey West

**Affiliations:** Integrated Mathematical Oncology Dept., H. Lee Moffitt Cancer Center & Research Institute, Tampa, FL; Dept. of Tumor Microenvironment and Metastasis, H. Lee Moffitt Cancer Center & Research Institute, Tampa, FL; Dept. of Biological Sciences, Indian Institute of Science Education and Research Kolkata; High School Internship Program, H. Lee Moffitt Cancer Center & Research Institute, Tampa, FL; Dept. of Head and Neck-Endocrine Oncology, H. Lee Moffitt Cancer Center & Research Institute, Tampa, FL

## Abstract

Direct observation of immune cell trafficking patterns and tumor-immune interactions is unlikely in human tumors with currently available technology, but computational simulations based on clinical data can provide insight to test hypotheses. It is hypothesized that patterns of collagen formation evolve as a mechanism of immune escape, but the exact nature of the interaction between immune cells and collagen is poorly understood. Spatial data quantifying the degree of collagen fiber alignment in squamous cell carcinomas indicates that late stage disease is associated with highly aligned fibers. Here, we introduce a computational modeling framework (called Lenia) to discriminate between two hypotheses: immune cell migration that moves 1) parallel or 2) perpendicular to collagen fiber orientation. The modeling recapitulates immune-ECM interactions where collagen patterns provide immune protection, leading to an emergent inverse relationship between disease stage and immune coverage. We also illustrate the capabilities of Lenia to model the evolution of tumor progression and immune predation. Lenia provides a flexible framework for considering a spectrum of local (cell-scale) to global (tumor-scale) dynamics by defining a kernel cell-cell interaction function that governs tumor growth dynamics under immune predation with immune cell migration. Mathematical modeling provides important mechanistic insights into cell interactions. Short-range interaction kernels provide a mechanism for tumor cell survival under conditions with strong Allee effects, while asymmetric tumor-immune interaction kernels lead to poor immune response. Thus, the length scale of tumor-immune interactions drives tumor growth and infiltration.

## 1 Introduction

LENIA (from the Latin *lenis* meaning “smooth”) is a cellular automata framework that allows for continuous space and time. Cellular automata are often used to study the “rules” that can recapitulate the collective behavior of interacting biological “agents” (e.g. individual cells or organisms). The development of Lenia was spurred by the search for rule-sets that lead to features that are important in artificial life: self-organization (morphogenesis) self-regulation (homeostasis), self-direction (motility), self-replication (reproduction), growth (development), response to stimuli or environmental factors, evolvability, adaptation^1,2^, and other emergent complexity^3^.

Many of these same features (e.g. cellular replication, growth, motility, evolvability, and adaptation) are important in the development of complex, multi-factorial diseases such as cancer. Thus, the purpose of this manuscript is to extend the Lenia framework as a tumor model, including immune predation and escape. We show how this is a natural extension, as Lenia has the capability to recapitulate 1) classical analytical models of cancer (ordinary differential equations), 2) common stochastic agent-based models of cancer, 3) standard models of multiple cell types competing on a Darwinian fitness landscapes (evolutionary game theory), and 4) generalized cell migration (e.g. chemotaxis models) common in cancer mathematical modeling literature.

### 1.1 The overlap in modeling artificial life and cancer evolution

In its original formulation, Lenia defines an on-lattice domain where each grid location can take a continuous density value. Methods of interactive evolutionary computation were used to find rule-sets that lead to defined, artificial “creatures” (see figure 1A for examples) which are capable of stable linear or angular motion or other life-like characteristics (figure 1B)^1,2^. These continuous creatures are reminiscent of the binary gliders, oscillators, and still-life creatures in Conway’s Game of Life^4^.

**Figure 1.**
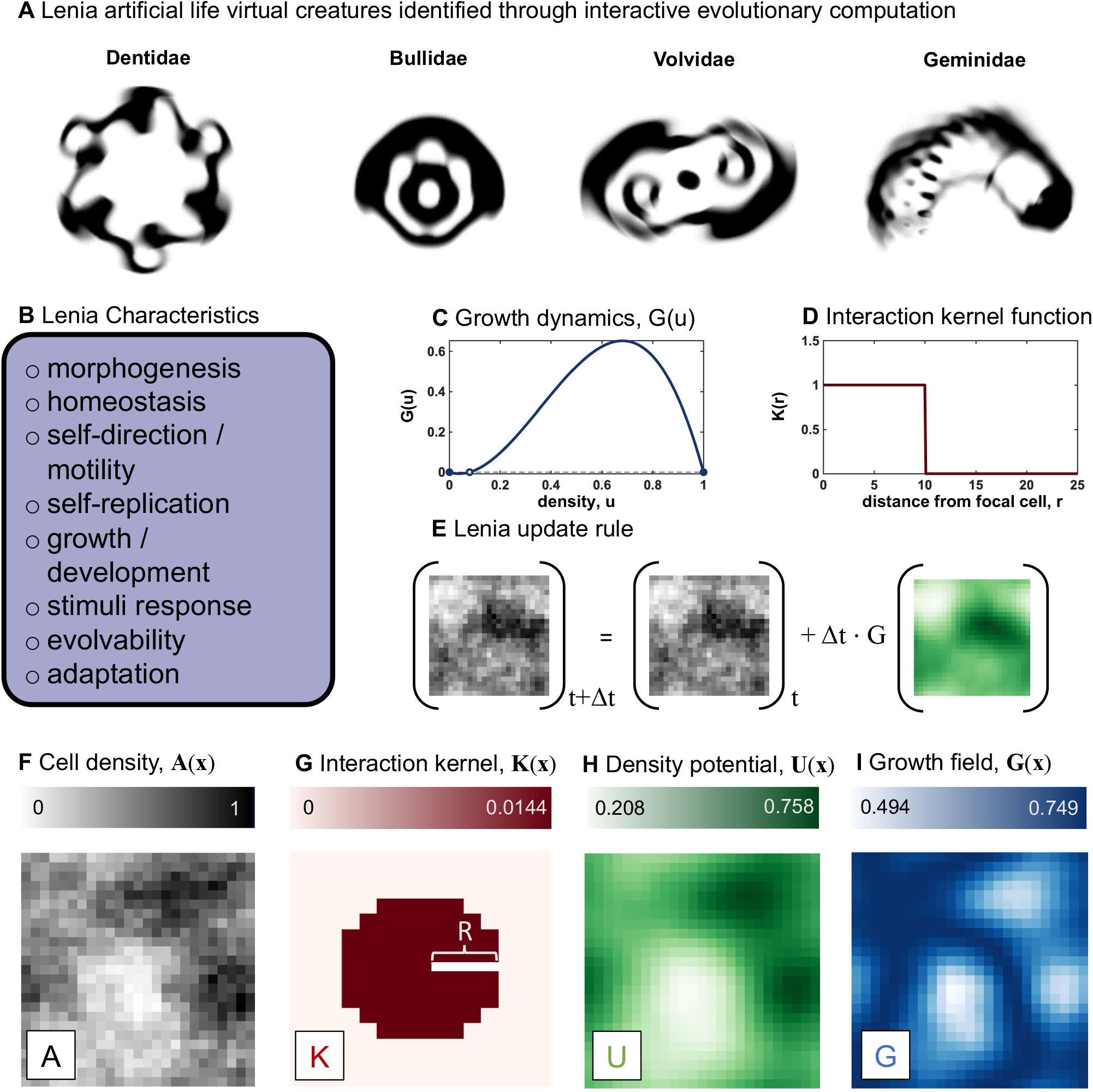
lenia as a cancer model: (a) lenia artificial life virtual creatures, reproduced from ref. 1. (b) list of characteristics possible to produce in lenia by varying the growth dynamics function (c) or the interaction kernel function (d). (e) adding a fraction of the growth field at each time step to the cell density, forms the lenia update rule, as seen in eqn. 1. (f) an example snapshot of a simulation shows the density of cells at each lattice location where the interaction kernel (g) specifies the nature of interaction of cells depending on their distance from each other. (h) the density potential, interpreted as a weighted average of interactions at each lattice location, is calculated as the convolution of **a**(**x**) and **k**(**x**). (i) the growth field is calculated by applying a growth map to the density potential.

Grid locations change according to an update rule describing the growth dynamics (figure 1C) paired with an interaction kernel function (figure 1D). For example, tumor cell density in a two-dimensional domain (figure 1E,F) is convolved with an interaction kernel (figure 1G) to calculate the “density potential” (figure 1H). Density potential is the surrounding density of grid locations weighted by kernel interaction function of radial distance from the focal cell. This defines a spatially-explicit growth field (figure 1I)^1^, whereby the update rule is enacted iteratively at each time step, t (figure 1E). Thus, a natural interpretation of Lenia as a cancer model is a spatial density map of cancer cells. Cancer cells grow according to a local density-dependent growth rate (e.g. logistic growth) which is a function of local cell density. Lenia can compare the effect of local (short-range kernels) versus global (long-range kernels) on growth dynamics.

There exists broad overlap in both methods and desired emergent behavior between models of artificial life and models of cancer growth (figure 1B). To adequately mimic biological realism in cancer, mathematical frameworks have implemented methods classified as 1) non-spatial or spatial, 2) deterministic or stochastic, and 3) single or multiple populations (cell types). Lenia allows for flexible model design to incorporate the full range of these classifications. Extensions to Lenia include Glaberish (a variant that leads to pattern formations that are stable in space or time^5^), Flow-Lenia (imposes mass conservation over time)^6^, Asymptotic-Lenia (asymptotic, smooth updating)^7^, Particle-Lenia (off-lattice)^8^, or a generalized reaction-diffusion implementation^9^.

Lenia is classified as a system of integro-differential equations (IDE)^10^. IDEs have been used to study cancer progression^11^ and treatment^12^ or to find approximations of stochastic ABMs in cancer^13^, but these formulations often focus on how interaction kernels alter the time scale of dynamics or the system stability. Lenia provides a unifying framework for direct comparison of well-mixed assumptions commonly used in ordinary differential equation (ODE) models (corresponding to Lenia’s long-range kernel) and local-scale competition commonly used in stochastic agent-based modeling (corresponding to Lenia’s short-range kernels). Next, we review common tumor-immune modeling frameworks of birth, death, and migration, and describe how Lenia may contribute to this body of literature.

### 1.2 Modeling spatial effects in tumor-immune dynamics

#### Spatio-temporal models of cell migration and the extracellular matrix

Immune cell migration is influenced by the spatial configuration of nearby collagen fibers^14, 15^. The density, orientation, and thickness of collagen fibers within the extracellular matrix (ECM) can influence migration through contact guidance thereby facilitating both migration and invasion^16^. Much of the mathematical modeling in this area focuses on recapitulating the spatial patterns of tumor-associated collagen signatures (TACS) which correlate with benign growth or invasive migration^17,18^. Models also aid in understanding the process of cancer cell metastasis as cells migrate through the ECM and extravasate into vasculature^19^.

#### Spatio-temporal models of cancer growth dynamics

Tumor growth can be expressed as a mathematical law using an ordinary differential equation (ODE)^20^ and used as a predictive tool in assessing the course of tumor progression^21, 22^. ODE frameworks use the well-mixed assumption, where each cell is equally likely to interact with each other cell in the tumor. Thus, there is a broad interest in understanding the effect of local, cell-scale contact inhibition on the emergent tumor-scale global growth dynamics^23–25^. The choice of spatial structure within a model (e.g. non-spatial, deme models, mesoscale, or boundary-driven growth) affects the evolutionary dynamics within heterogeneous tumors^26–29^, tumor growth dynamics^30, 31^ and the emergent pattern formation^32–34^.

#### Spatio-temporal models of tumor-immune interactions

Tumors do not grow in isolation but must develop immune escape mechanisms to avoid extinction. Tumor-immune interactions have been described using logistic tumor growth with immune recruitment and exhaustion^35^ or as a predator-prey relationship^36^. Immune predation has likewise been modelled using integro-differential equations^37–40^, mesoscopic models^41^, or stochastic agent-based methods^42^.

Importantly, classical predator-prey models may be insufficient to describe tumor-immune dynamics without also considering mechanisms for avoiding oscillatory behavior common to these models^43^. Thus, in the models developed herein we include Allee effects, whereby tumors require a cooperative benefit of close aggregation when at lower densities^44, 45^. Allee effects lead to a decreased growth benefit when tumor density is low, and thus are proposed to enable immune predation to fully eliminate tumors in the absence of immune escape^43^.

#### Lenia as a birth-death-migration model

In summary, stochastic birth-death-migration models are commonly used to model tumor-immune growth dynamics, often with a focus on deriving approximations that map local interaction dynamics to global mean-field approximations. Alterations in cell migration, proliferation, and death parameters may lead to spatial heterogeneity in cell density, reducing the accuracy of naive mean-field approximations. Thus, it is convenient for a cancer modeling framework to include 1) non-spatial or spatial, 2) deterministic or stochastic, and 3) single or multiple populations (cell types). Below, we begin by implementing a data-driven Lenia model of immune infiltration in head and neck squamous cell carcinomas using Particle (off-lattice) Lenia. Next, we illustrate how classical growth dynamics models (e.g. logistic growth) are a special case of Lenia (without immune predation). Finally, we illustrate how multi-channel Lenia can be used to model cell-cell (tumor-immune) spatial interactions.

## 2 Results

### 2.1 Stage-dependent collagen alignment predicts poor immune infiltration

Direct observation of immune cell trafficking patterns is unlikely in clinical tumors, but computational simulations based on clinical data can provide insight to test hypotheses. We begin by developing a model of immune cell infiltration using an extension of Particle Lenia^8^. Figure 2A introduces the image analysis to computational model pipeline. Second harmonic generation (SHG) is performed on a cohort of HPV-negative head and neck squamous cell carcinomas (HNSCC) to quantify collagen density, **D**(**x**). Each patient has a selected region of interest (ROI) to perform SHG that are subsequently analyzed using OrientationJ ImageJ plugin to determine fiber alignment,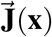^46^. Each pixel is color-coded according to the directionality of collagen fibers, such that fibers aligned in a single direction are colored with the same hue on the red-blue-green-yellow spectrum (figure 2A, semi-circle legend color-bar). The microenvironment gradient field, ∇**M**(**x**) at each grid location **x** is determined by the convolution of interaction kernel and the OrientationJ vector field, weighted by collagen density (see Methods). Thus, immune cells move along the microenvironment gradient field in the same direction as collagen fiber alignment. In Lenia, immune cell movement direction is determined by the average collagen fiber alignment within the immune cell’s interaction kernel radius, *R* (see Methods; eqn. 8).

**Figure 2.**
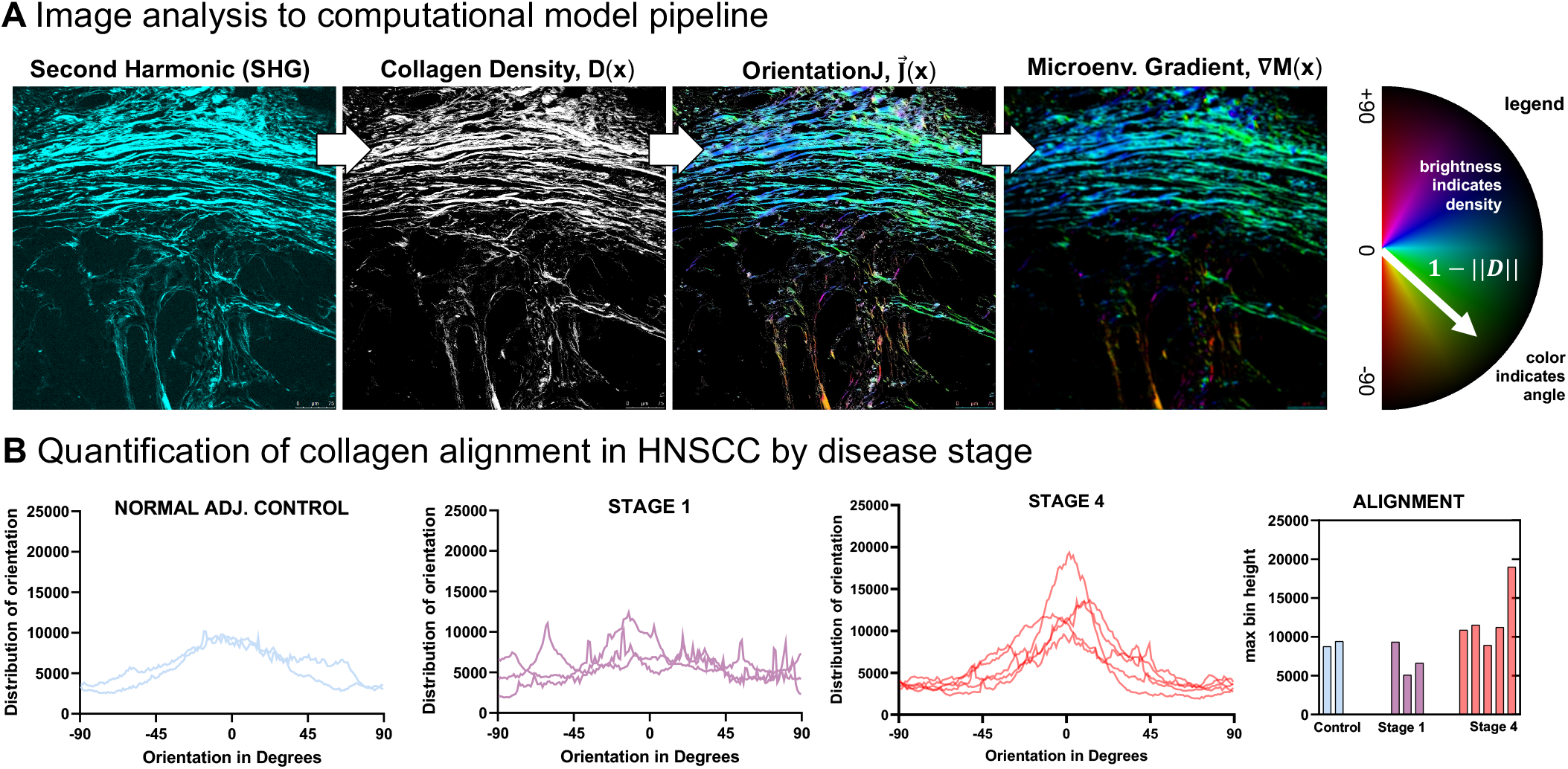
Collagen alignment in HNSCC. (A) Pipeline of image analysis to computational model using second harmonic generation to determine collagen density and subsequently the alignment and microenvironmental gradient. (B) Alignment collagen fibers is determined using OrientationJ ImageJ plugin^46^; alignment tends to increase by disease stage.

#### Quantification of collagen alignment by disease stage

Late-stage HNSCC are associated with increased deposition and alignment of fibrillar collagens^47^. We hypothesize that poor outcomes are partially due to poor immune cell infiltration caused by deposition of a dense collagen barrier. Imaging data are quantified by stage in figure 2B, which illustrates the correlation of disease stage with collagen density and alignment. Normal adjacent control tissue and early stage HNSCC’s are associated with sparse and heterogeneous collagen alignment, while late-stage HNSCC tends to contain highly aligned collagen fiber patches. We hypothesized that late-stage disease is associated with collagen structures which lead to poor immune infiltration. In the next section, we next employed Lenia to test two plausible cell migration models describing the movement of immune cells in response to collagen: parallel or perpendicular movement.

#### Simulated effect of collagen alignment on immune surveillance

Within the immune cell trafficking model each simulated immune cell moves along a microenvironment gradient field determined by collagen fiber alignment (Methods; eqn. 8). We consider two hypothesized models of the influence of collagen fiber alignment on immune cell trafficking: perpendicular or parallel to collagen fibers (figure 3A). Immune cells are initially evenly distributed on the edge of the domain, and we quantify the area of surveillance covered (simplified example shown for six immune cells per side in figure 3B). The total simulated time is chosen such that immune cells in an empty domain would migrate in a linear, unaltered fashion and provide 100% coverage of the domain.

**Figure 3.**
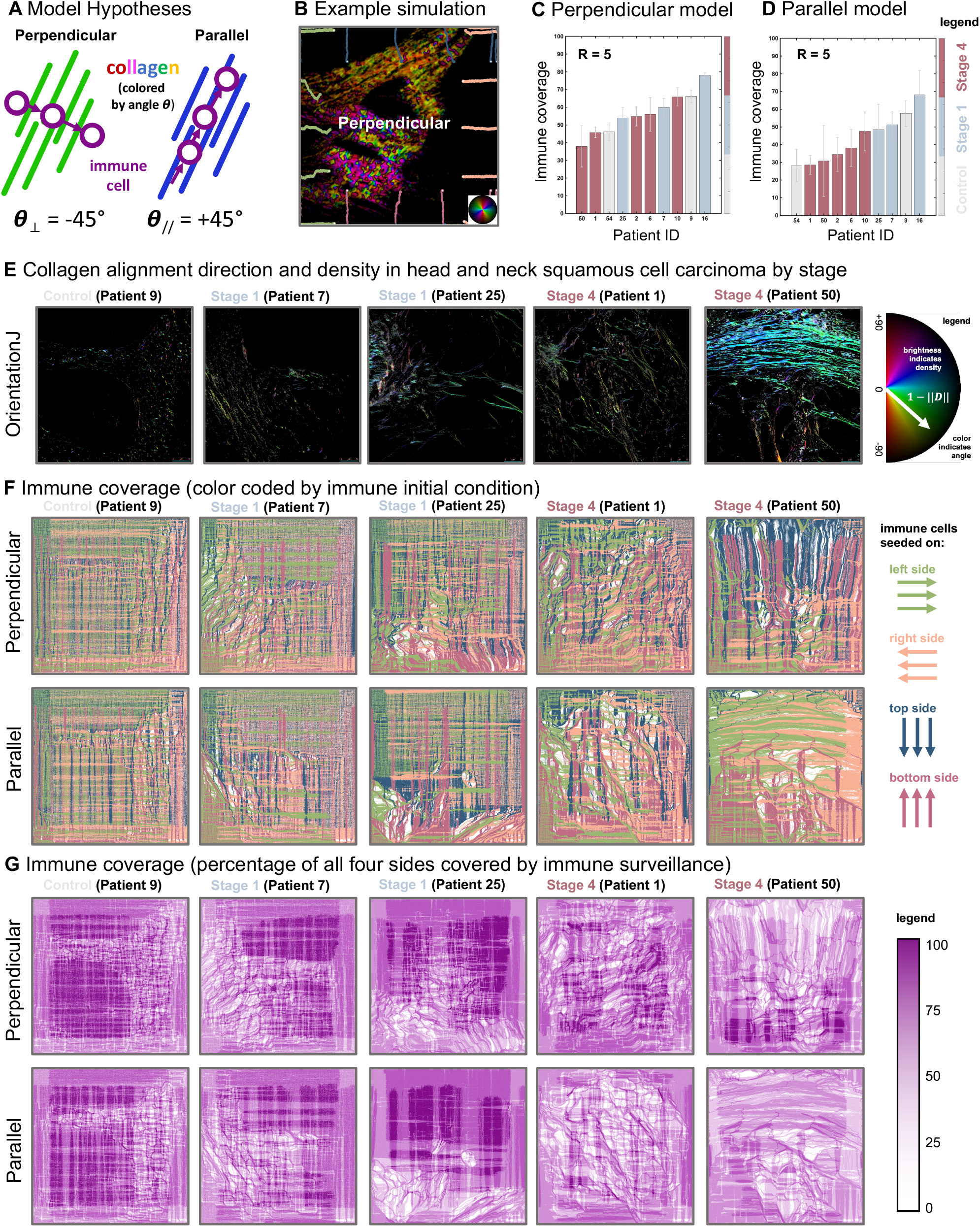
Immune cell trafficking model in Particle Lenia. (A) Hypothesized models of the influence of collagen fiber alignment on immune cell trafficking. See corresponding Supplementary Video S1. (B) example simulation showing immune cell infiltration (circles) with track indicating path taken. Cells seed on all four sides, color-coded by initial side (see panel F). (C, D) Simulated immune coverage for perpendicular (C) or parallel (D) immune trafficking model, colored by stage. (E) Representative sample ROI alignment and density of collagen fibers (using OrientationJ ImageJ plugin^46^) ordered by disease stage. (F) Immune coverage, color-coded by immune initial condition of which of the four sides: left, right, top, bottom (see legend). The color of each pixel is determined the most trafficked side. (G) Immune coverage as the percentage of all four sides covered by immune surveillance. For example, if no immune cells reached this pixel it is white, if all four sides reached this pixel, it is dark purple. See corresponding Supplementary Video S2.

Results indicate an overall higher level of immune coverage in the perpendicular model than in parallel (figure 3C and D, respectively). Interestingly, the parallel model shows a stronger anti-correlation between immune coverage and disease stage. All stage 4 patients rank lower in immune coverage compared to patients with stage 1 tumors (figure 3D). Of note, it’s difficult to interpret immune coverage for control tissues as the absence of immune infiltrates may result from lower collagen density typical of normal tissue and that is not well-aligned, or it could simply be due to absence of diseased tissue (i.e., tumor) and thus the absence of signaling cues required to induce immune cell trafficking. Late-stage HNSCC tumors are associated with a more hypoxia-related signature characterized by a low ratio of CD8+ T cells to FOXP3+ regulatory T cells^48^. Thus, as the parallel model consistently predicts lower immune coverage for late-stage tumors, this model may be more biologically relevant. This also supports previous experimental findings that suggest immune cells move parallel to collagen structures^49^. Studies have shown that collagen fibers are often perpendicular to the outer boundary of tumors^50, 51^, suggesting that a parallel model of immune migration would reduce immune infiltration into the tumor. This finding supports the notion of the parallel immune migration model as a method of immune escape. Note: we do not consider variable immune response here (which may vary with disease stage) as the total number of immune cells seeded is constant across all patient simulations.

Individual model simulations are shown for a model seeded from the corresponding collagen ROI in figure 3E. Immune cells are seeded along each edge (left, right, top, bottom) of the domain (*N* = 5 cells per pixel; figure 3F). Typically, as disease stage increases (left to right), immune cell coverage decreases as seen in figure 3G. We define immune coverage as the percentage of the domain covered (100% corresponds to pixels which are covered by at least one immune cell trafficked from each of the four sides). Well-aligned collagen structures (e.g. stage 4) result in a strongly directed migration, deviating immune cells from providing a well-mixed coverage of the domain.

### 2.2 Increased range of interaction increases tumor diffusivity and speeds growth

The Particle Lenia modeling above provides insight into immune migration behavior but neglects tumor growth. In the following sections, we implement a classic Lenia model of tumor (figures 4 and figure 5) and immune (figure 6) dynamics to illustrate the effect of the range of interaction kernel on spatial interactions. First, we implement a single population (one cell type) model of tumor growth dynamics in figure 4.

**Figure 4.**
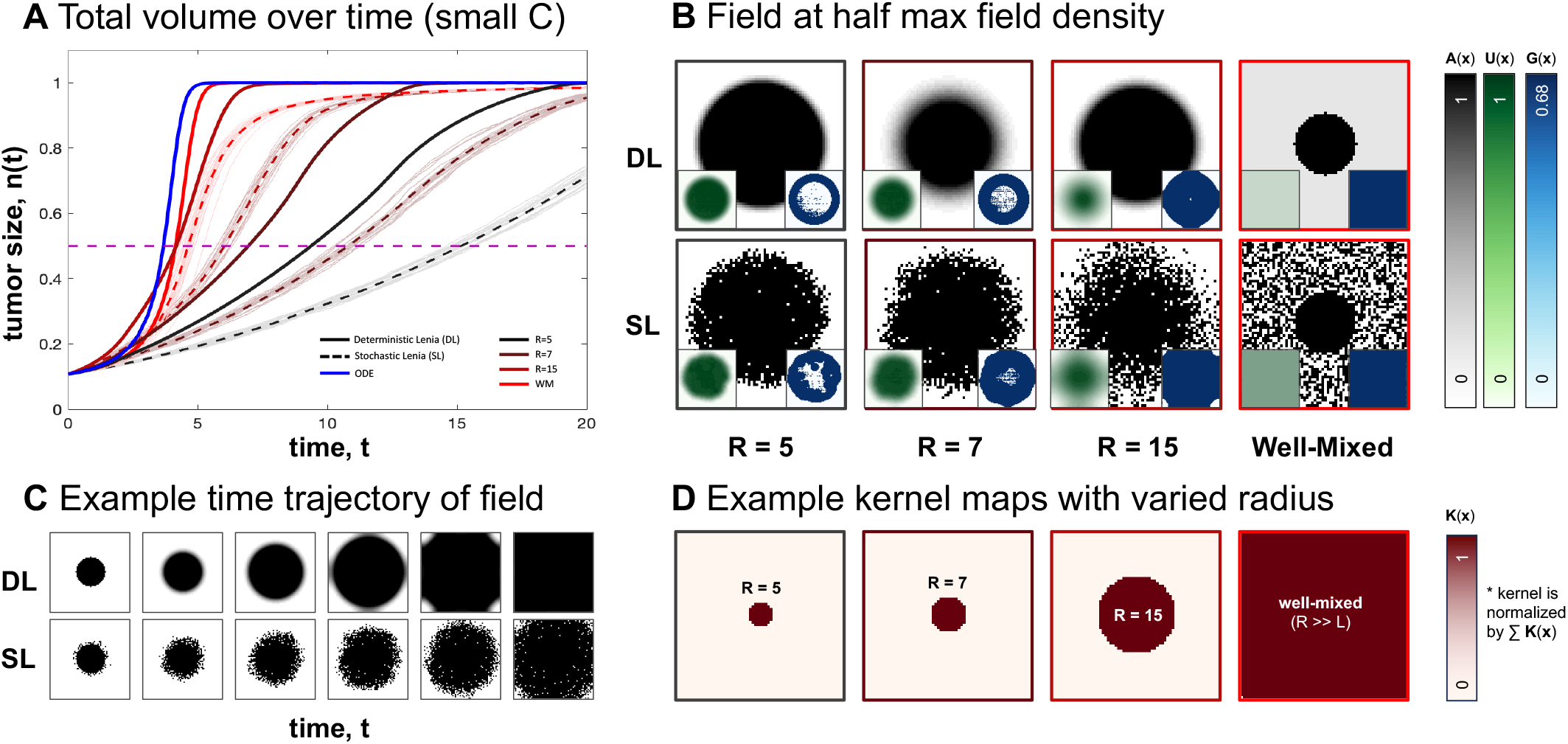
Cancer growth dynamics in Lenia. (A) Average cell density over time for varying kernel sizes, for deterministic and stochastic Lenia. Averages over stochastic runs are shown in dotted lines and individual trajectories are shown in translucent lines. ODE solution (shown in blue) matches simulation with a well-mixed kernel. (B) State of field at half maximal capacity for different kernel sizes for Stochastic Lenia (SL) or Deterministic Lenia (DL). Density potential and Growth distribution shown in inset. (C) State of field at progressive time points for kernel size 4 for deterministic and stochastic Lenia. (D) Kernels used in models in figures (B).

**Figure 5.**
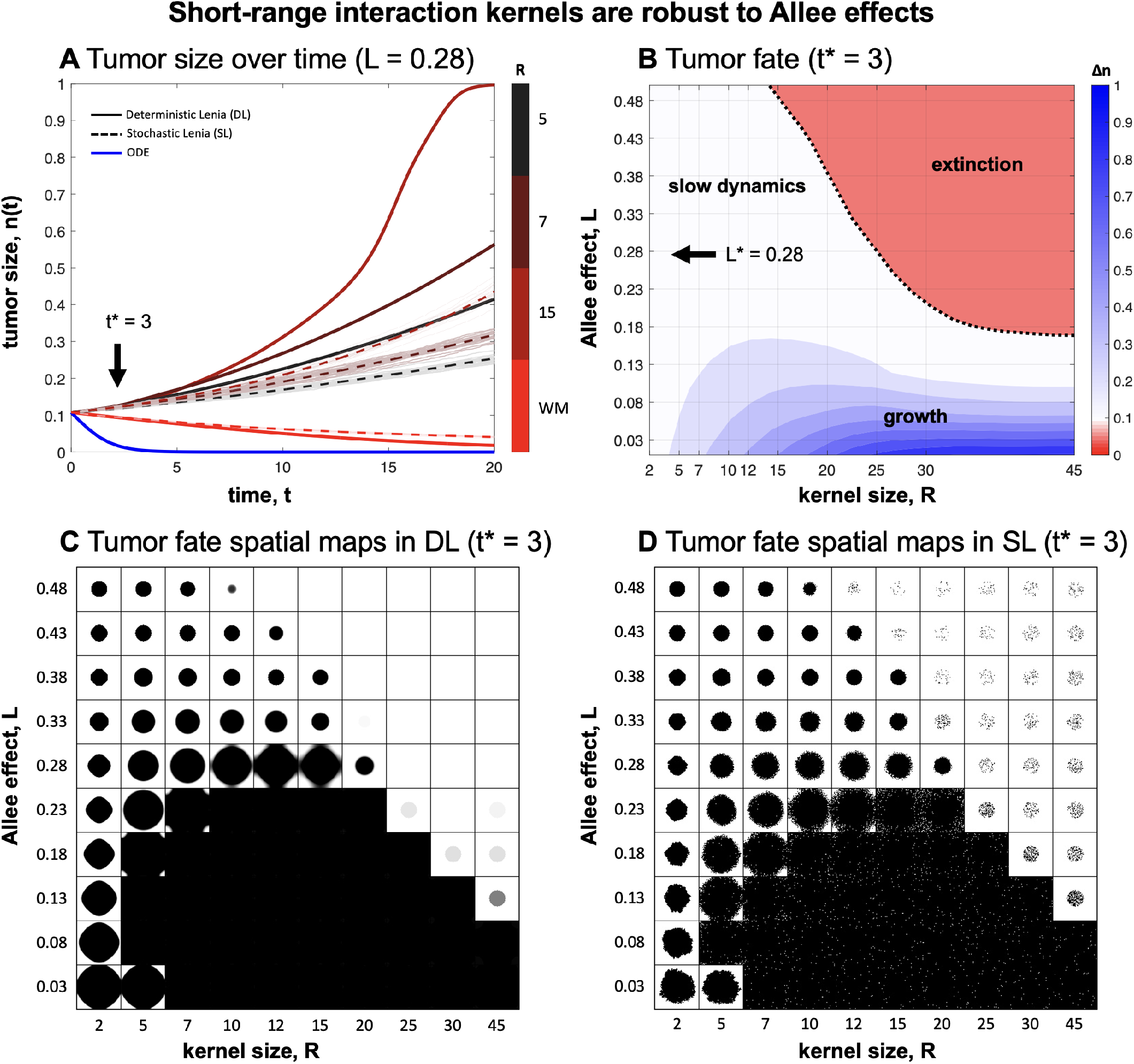
Short-range interaction kernels are more robust to Allee effects. (A) Average cell density over time for varying kernel sizes, for deterministic and stochastic Lenia. Averages over stochastic runs are shown in dotted lines and individual trajectories are shown in translucent lines. ODE solution shown in blue. (B) Long-term tumor fate for deterministic Lenia where trajectories that eventually result in extinction are shown in red, or growth shown in blue (colorbar indicates change in tumor size at *t*^*∗*^ = 3. (C, D) Spatial maps shown for deterministic Lenia (C) and stochastic Lenia (D) at time *t*^*∗*^ = 3. See corresponding Supplementary Video S3.

**Figure 6.**
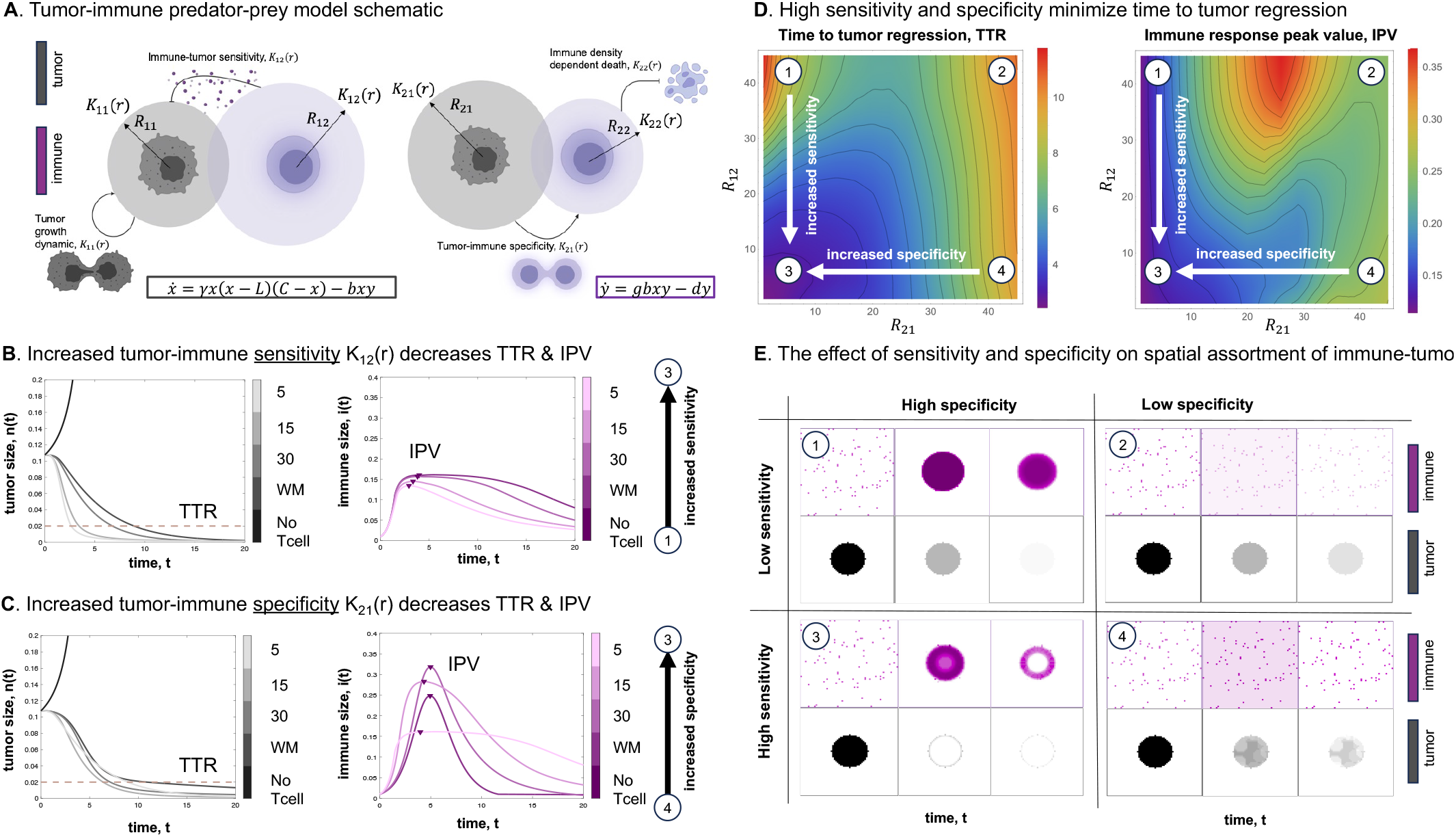
Competition dynamics in Lenia. **(A)** Schematic representation of the tumor-immune predator-prey model, illustrating various kernel sizes for tumor growth (*K*_11_), predation distance (*K*_12_), immune recruitment (*K*_21_), and density-dependent death distance in the immune system (*K*_22_). **(B, C)** Impact of different predation distances (*K*_12_) and immune recruitment (*K*_21_) on the integration results of the tumor-immune predator-prey model, with other kernels held well-mixed. **(D)** Effect of varying predation and recruitment (*K*_12_ and *K*_21_) on the time to tumor regression (considered when there are 2% tumor cells in the area) and the peak value in the immune response. **(E)** Spatial outcomes of the tumor and immune response in the model, showcasing the influence of different kernel sizes. See corresponding Supplementary Video S4. Unless otherwise noted, parameters used are *γ* = 5, *b* = 12, *g* = 1.5, *d* = 1, *L* = 0.08 (see eqns. 6-7).

Growth dynamics are described by logistic growth with an Allee threshold (Methods, eqn. 4), such that tumors seeded above the Allee threshold have positive growth. The solution to this ordinary differential equation is shown in blue in figure 4A. Comparing deterministic Lenia (figure 4A; solid lines) illustrates that systematically reducing the effective range of the interaction kernel reduces the rate of growth, and alters the temporal dynamics. Smaller interaction kernels (e.g. black lines) are characterized by slow growth due to the presence of contact inhibition in small interaction neighborhoods.

Reduction in kernel size (different kernel sizes used in figure 4B are shown in figure 4D) also alters the spatial variegation of tumor density. For example, short-range kernels (figure 4B, left) are characterized by sharp, well-defined boundary drive growth while long-range kernels (figure 4B, right) are associated with diffuse patterns.

This pattern holds when considering either a deterministic (figure 4B; Deterministic Lenia, DL) or a stochastic (figure 4B; Stochastic Lenia, SL) update rule (see Supplemental section S7.2). The average of many stochastic realizations is shown in dashed lines (figure 4A), which tend to underestimate the deterministic simulation with the corresponding kernel size. This is due in part to the fact that the cell density map **A**(**x**) in stochastic Lenia is restricted to integer values only, which is common practice in standard agent-based models (compare figure 4B, where only values of 0 or 1 are allowable). These results illustrate the utility of Lenia to interpolate between two standard modeling approaches: 1) ordinary differential equations and 2) stochastic, local agent-based methods. Lenia extends standard methods to allow investigation of arbitrary interaction kernels.

### 2.3 Short-range interaction kernels are more robust to Allee effects

Next, we investigate the interplay between the Allee threshold and the interaction kernel. Logistic growth with an Allee threshold has an unstable equilibrium which drives the population to extinction when tumor density is below the threshold. As shown in figure 5A, simulating logistic growth with Allee in an ODE (blue line) or a well-mixed (red line; WM) both result in eventual extinction. Interestingly, reducing the interaction kernel distance, R, results in positive tumor growth (e.g. black line). This effect occurs for a range of Allee threshold values and kernel sizes, shown in figure 5B. This clearly illustrates an expanded region of positive growth when kernel size is low, across a wide range of threshold values. Smaller neighborhoods increase the likelihood that a single cell’s neighborhood will maintain a sufficient density above the Allee threshold, and thus maintain positive growth.

While stochastic Lenia is typically associated with slower temporal scales (figure 5A, dashed lines), the long-term tumor fate (extinction or growth) are analogous in both models and spatial variegation patterns are also similar (figure 5C,D). These results suggest that well-mixed model formulation may overestimate the role of an Allee effect (thus overestimating the likelihood tumor extinction). In the next section, we investigate the effect of interaction kernel size on immune predation of tumors.

### 2.4 High immune specificity and sensitivity maximize tumor regression

In order to investigate the effect of short-range and long-range interactions between tumor and immune cells, we employ the multi-channel Lenia model to simulate the dynamics of in a tumor-immune (figure 6). Tumor-immune dynamics are described by the predator-prey model in eqns. 6 and 7. Similar to the previous figure, this model contains an Allee threshold whereby tumor cells are driven to extinction below this threshold even in the absence of immune cells^43^.

We consider four interaction kernels: tumor-tumor, tumor-immune, immune-tumor, and immune-immune (figure 6A). We define the tumor-immune kernel (*K*_12_) as the **immune sensitivity**, whereby a short-range *K*_12_ is characterized by an increase in immune kill rate and decreasing the time to tumor regression (TTR) as seen in figure 6B. To gain intuition, one can consider the tumor-immune kernel as representing the average distance traveled by an immune cell per unit of tumor kill. For example, a large tumor-immune kernel may be represent a tumor with high PD-L1 expression, requiring immune cells to travel a further distance before effective immune activity.

Next, we define the immune-tumor kernel as the **immune specificity**, whereby a short-range *K*_21_ leads to more localized recruitment of immune cells in close proximity to tumor cells. A reduction in the recruitment distance (small *K*_21_) leads to the production of an immune response in closer proximity to tumor cells (refer to figure 6E). To gain intuition about the immune-tumor kernel, one can consider this kernel as representing the average distance an immune cell is recruited by a tumor. An immunologically quiet tumor is represented by a large kernel with diffuse, undirected recruitment, while increased tumor antigenicity is represented by a small kernel with strong, localized recruitment. The resulting immune peak value, IPV (figures 6B, C) depends on both **immune sensitivity** and **immune specificity**.

As seen in figure 6D, both immune specificity and immune sensitivity are required to maximize tumor regression (case 3). This effect is illustrated by observing spatial maps over time in figure 6E. High specificity results in rapid recruitment concentrated within the tumor, but these immune cells are most effective at removing tumor when sensitivity is high (case 3). In contrast, low specificity results in a diffuse spatial pattern of immune recruitment through the domain, where most immune cells are not co-localized with tumor cells, effectively wasting a majority of immune response (cases 2 and 4). Consequently, immune cells that do infiltrate tumor are rare but strongly effective, leading to a heterogeneous tumor kill (case 4). The effect of asymmetric interaction kernels leading to differences in spatial variegation and patterning in tumor and immune cells is shown for a range of sensitivity and specificity values in supplemental figure S1.

## 3 Discussion

Given the fact that direct observation of immune cell migration behaviors is limited with currently available methods, computational modeling has emerged as a viable method for interrogating tumor-immune interactions. Tumors under immune predation may alter their surrounding extracellular matrix (figure 2), leading to patterns of collagen formation as a mechanism of immune escape. Here, our modeling framework in Lenia is able to distinguish between parallel and perpendicular modes of immune cell migration as a function of collagen fiber alignment (figure 3). Results indicate that a parallel mode of migration is more likely, given the inverse correlation between model-predicted immune infiltration and disease stage. Previously, aligned collagen fibers have been observed perpendicular to the tumor periphery^50, 51^, lending supporting evidence to a parallel model of immune migration that reduces immune infiltration. Despite neglecting alternative mechanisms of immune escape (e.g. acid-inactivation, immune checkpoint expression, or immune exhaustion), the immune infiltration model shows an inverse relationship between disease stage and immune coverage as an emergent phenomenon (figure 3).

Methods in artificial life represent a promising cross-over potential in cancer modeling. For example, Lenia demonstrates the capability to recapitulate important features required for modeling the evolution and ecology of cancer: growth dynamics (deterministic/stochastic), cell-cell interactions, and cell migration. To our knowledge, we have introduced the first example of Lenia applied as a cancer model^52^. While here we focus on modeling tumor-immune spatial interactions, the role of spatial structure more broadly has important implications for optimizing cancer treatment^53, 54^, modulating evolutionary dynamics of tumor heterogeneity^27, 29, 55^, and altering cell-cell competitive dynamics^56^ even when using the same interaction rules^57, 58^. Lenia is flexible enough to mimic non-spatial (well-mixed) ordinary differential equations (with sufficiently large kernel sizes) as well as small-scale spatial neighborhoods commonly used in agent-based methods (e.g. Moore or von Neumann neighborhoods). It is also straightforward to simulate deterministic dynamics and the corresponding stochastic dynamics. Additionally, Lenia can simulate ecological interactions between multiple cell types in heterogeneous tumors. We have shown the ability of Lenia to recapitulate common models of tumor-immune dynamics (logistic-Allee). Finally, we illustrate the ability of Particle Lenia to recapitulate cell migration and trafficking in an off-lattice, with an application in squamous cell carcinomas. However, Lenia provides several natural advantages over traditional birth-death-migration models. Kernel convolution can be performed using fast-Fourier transforms, drastically decreasing the computational cost of simulations. Additionally, convolution of extrinsic migration gradient can be pre-computed, increasing computational efficiency.

In addition to flexible methodology, Lenia has shown promise in understanding the mechanics of tumor-immune interactions. We have illustrated that immune-ECM interactions decrease immune infiltration (figure 3). Next, we show that short-range interaction kernels provide a mechanism for tumor cell survival under conditions for strong Allee effects (figure 5). We have also shown the importance of the length scales of immune sensitivity and specificity in immune response (figure 6). In the absence of strong sensitivity or specificity poor immune response leads to prolonged tumor growth and in some cases, immune escape. Differential interaction kernels alter the spatial patterns of tumor-immune cells (figure S1), leading us to hypothesize the possibility of making inferences of kernel sizes based on spatial co-localization of cells in future work. Here, we may draw inspiration from literature in ecology where kernel inferences are drawn from plant seed dispersal patterns^59^. Thus, in each of these examples, the characteristic kernel distance of tumor-immune-ECM interactions drives tumor growth and immune infiltration.

### Methods

#### 4.1 Lenia Cellular Automata

The notation here follows ref. 1, which we re-state here for clarity. Lenia is a cellular automata defined by five components: 1) a discrete grid of points, *ℒ*, (herein we consider two-dimensional lattice grid *ℒ* ∈ ℝ^2^), 2) the range of allowable states *𝒮* = [*A*_min_, *A*_max_], 3) the local update rule *φ* : *𝒮* ^*𝒩*^ *→ 𝒮* is the local update rule, 4) the neighborhood around each focal cell at location **x**, given by *𝒩*_**x**_ = {**x** + **n** : **n** ∈ 𝒥}, and 5) time steps *t* ∈ *𝒯*. The spatial *cell density* map configuration is defined by **A**^*t*^ : *ℒ → 𝒮* : the collection of states over the whole grid, at time *t* ∈ 𝒥, the timeline. Herein, we let **A** be a two-dimensional matrix where **A**(**x**) represents the cell density at grid location **x** which is updated via the following equation:

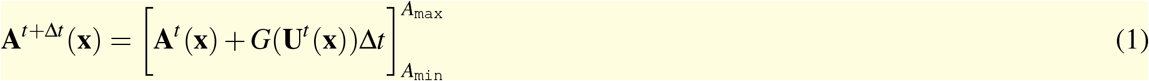

where *G*(*u*) is the density-dependent growth function defined on the interval *u*∈ [*A*_min_, *A*_max_]. For each point **x**, the convolution of a kernel **K** with **A**^*t*^ (*𝒩*_**x**_) is calculated to compute the potential distribution **U**^*t*^ :

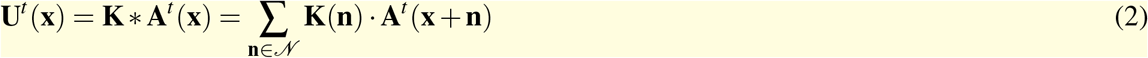

Thus, the growth is defined at every lattice location: **G**^*t*^ (**x**) := *G*(**U**^*t*^ (**x**)). A fraction of the growth distribution is added to each site at each time step (Euler approximation update rule) and the value at the site is clipped to [*A*_min_, *A*_max_]. We implement the computational model in the Hybrid Automata Library framework (Java)^60^.

#### 4.2 Interpretation of Lenia as a cancer model

Next, we make explicit the connection to cancer modeling. Lenia provides a natural framework for constructing mathematical models of cancer cell growth, competition, and migration, where each lattice location **x** contains a density of tumor cells, **A**^*t*^ (**x**). When modeling tumor dynamics, set *A*_min_ = 0 and *A*_max_ = *C* where *C* represents the maximum resource- or space-limited carrying capacity of cells within each single lattice location. Classic cell interaction neighborhoods such as von Neumann (the four nearest neighbors; *R* = 1) or Moore (eight nearest neighbors; *R* = 1.8) can be written as an interaction kernel function, defined as a circle of radius *R*:

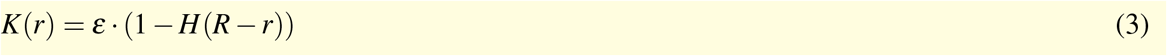

where *H* is the Heaviside function, and *ε* is chosen to normalize the elements of the kernel to sum to one. Thus, **U**(**x**) represents a potential density that is perceived by the focal location, determining the location-specific growth rate of cells, *G*(**U**(**x**)). The density potential is the convolution of cell density within the neighborhood, **A**(*𝒩*_*x*_), weighted (convolved) by the values in the kernel function. Typical kernel functions in cancer modeling are likely to be non-increasing functions of the radial distance from the focal location. For notational convenience, define the mean density of the field as 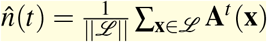, where ||*ℒ*||is the total number of lattice locations in domain *ℒ*.

Lenia can be extended to a stochastic setting by restricting only integer values on each lattice point (**A**(**x**) ∈ [0, 1, 2, …, *A*_max_]) with stochastic updates (see Supplemental Section S7.2). We note that classic agent-based methods often restrict each lattice location to contain at most a single cell (*A*_max_ = 1) and make the implicit assumption that cells divide into surrounding empty locations with probability *b*. This is qualitatively similar to defining a Moore-like or von Neumann-like interaction kernel with *G*(**U**(**x**)) = *b*.

##### 4.2.1 Tumor growth dynamics model

We first model tumor growth as a local logistic growth dynamic, shown in figure 1E, and F given by:

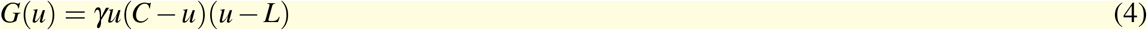

where *γ* is the intrinsic growth rate of tumor cells, *C* is the carrying capacity (maximum density) of each lattice location, L is the allee effect, and *u* is the density potential (eqn. 2).

##### 4.2.2 Tumor-immune predator-prey model

Using multi-channel Lenia, we can extend the model to account for cell density of multiple cell types, **A**_*i*_(**x**) where *i* ∈ {1, 2, …, *N}*. For example, herein we consider competition between *N* = 2 cell types that each have a growth field which is a function of the density potential for both cell types: 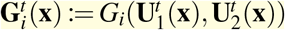.

In this two-type case, the dynamics are given by:

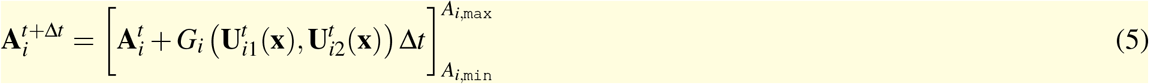

Thus, the tumor-immune model can be written as a system of two growth functions:

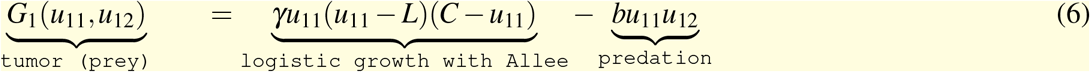

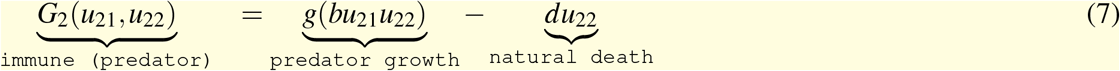

where *u*_*ij*_ represents the density of cell type *j* perceived by cell type *i*. The Allee threshold is given by *L*, carrying capacity of tumor cells is *C*, and predation rate is *b*. Immune cells and expand at rate *g* (weighted by immune predation response) and die at rate *d*.

###### Modeling cell migration using Particle Lenia

A recent extension referred to as Particle Lenia extends the mathematical framework to off-lattice. The location of each cell *i* is given by vector **p**_*i*_ := {*x, y*} corresponding to its location in two-dimensions. Movement of the particle is determined by the local gradients of the energy and chemotactic migration fields: **E**(**p**_*i*_) and **M**(**p**_*i*_), respectively.

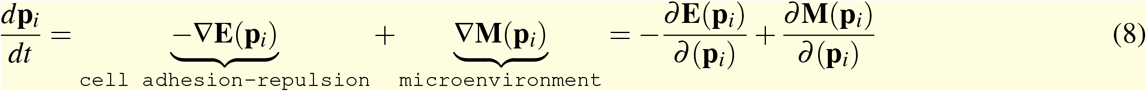

Particles (i.e. cells) in this model move against the local gradient of the energy field, representing particle interactions^*a*^. The cell-cell repulsion and adhesion fields are a function of the potential field, **E**^*t*^ (**x**) = *E*(**U**^*t*^ (**x**)) (see Supplemental info section S7.3). Similarly, an extrinsic microenvironmental field **M**(**p**_*i*_), defines the chemotactic direction of cells moving toward a nutrient-based or bio-mechanical gradient. The cell position update rule can be deterministic (following the highest point of gradient) or stochastic (weighted in the direction of gradient).

###### Modeling the effect of collagen on T-cell migration

In this manuscript, we assume negligible T-cell adhesive-repulsive interactions such that *E*(**U**^*t*^ (**x**)) = 0. Instead, T-cells move in a directed fashion influenced by the surrounding extrinsic microenvironment according to the following equation:

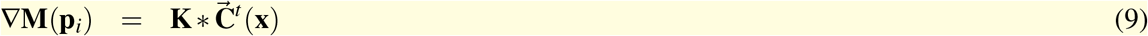

where **K** is the cell-microenvironment interaction kernel (see eqn. 3), 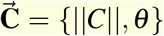 is a vector field determined by collagen fiber alignment. At each time point, noise is added to the collagen vector field’s angle such that *θ ′* = *θ* + *R*(*σ*) where *R* is a uniform random variable on [0, *σ* ] radians.

In this way, we model the influence of collagen fibers to direct T-cell movement as a weighted vector summation of the collagen fiber alignment in the focal T-cell’s local neighborhood. The collagen fiber alignment vector field is inversely proportional to the density of collagen (cells travel slowly in dense fibers). We consider two plausible models of the influence of collagen on T-cell migration: T-cells travel 1) in parallel to collagen fibers or 2) perpendicular to collagen fibers, denoted ∥ and ⊥, respectively.

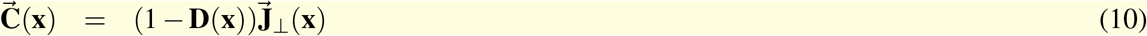

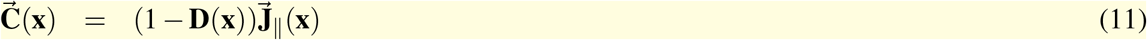

where 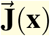 is collagen fiber alignment (a unit vector) derived from clinical imaging, detailed in the next section below, scaled by the inverse of the normalized density of collagen at each grid location, **D**(**x**) ∈ [0, 1].

###### Immune cell migration field defined using OrientationJ

Tumor microarray (TMA) having FFPE tissue samples from HPV-negative HNSCC patient cohort from the H. Lee Moffitt Cancer Center and Research Institute (IRB: MCC#18754) was analyzed to determine collagen alignment in the tumor microenvironment. The TMA cores were imaged using a Leica SP5 AOBS multiphoton confocal microscope and a 25x, HC IR Apo, NA 0.95 water immersion objective. The excitation wavelength was tuned to 880 nm and emitted light was detected at 440 nm using bandpass filters for detecting the SHG signal of collagen. The SHG images were analyzed using the Orientation J plugin^46^ in ImageJ software which provides both visual and quantitative data outputs for collagen fiber alignment^62^.

## 5 Acknowledgments

Authors gratefully acknowledge funding support from the National Cancer Institute via the Cancer Systems Biology Consortium U54CA274507 (A. Anderson), Florida Biomedical Research Program James & Esther King grant (21K04) (C. Chung), the Moffitt Cancer Center’s “Center of Excellence for Evolutionary Therapy” (J. West), the Moffitt Cancer Center’s “Cancer Biology & Evolution Program” pilot award (A. Amelio & J. West). The authors also would like to thank support from the staff in Moffitt’s Analytical Microscopy and Quantitative Imaging Cores: Joe Johnson and Abdalah Mahmoud. The Analytical Microscopy and Quantitative Imaging Cores are supported in part by the NCI Center Core Support Grant P30-CA076292 to the H. Lee Moffitt Comprehensive Cancer Center.

## 6 Code Availability

Code for an implementation of Lenia in the Hybrid Automata Library^60^ framework (Java) can be found here: https://github.com/mathonco/lenia-in-hal^52^.

## S7 Supplemental Information

### S7.1 Spatial variegation of tumor-immune interactions

We assess the degree of spatial variegation and patterning in tumor and immune cells as a function of sensitivity and specificity in figure S1. Spatial maps are shown at time to tumor regression (TTR; see figure 6), and thus all tumors have identical total size in figure S1A. Differences in spatial patterns of tumor density are caused by differential immune recruitment (specificity) and strength of predation (sensitivity). As the immune specificity narrows (*r*_21_ reduces to smaller values), immune cell concentration strongly correlates with the tumor’s location with a high degree of specificity (figure S1B, right-to-left). This specificity has only a marginal effect on tumors if the range of sensitivity is diffuse (large *r*_12_ values). As sensitivity narrows, immune predation is more highly targeted and effective, killing tumor cells, especially in the core (figure S1A, bottom-left). A ring-shaped tumor is left, due to proliferating cells on the rim.

**Figure S1.**
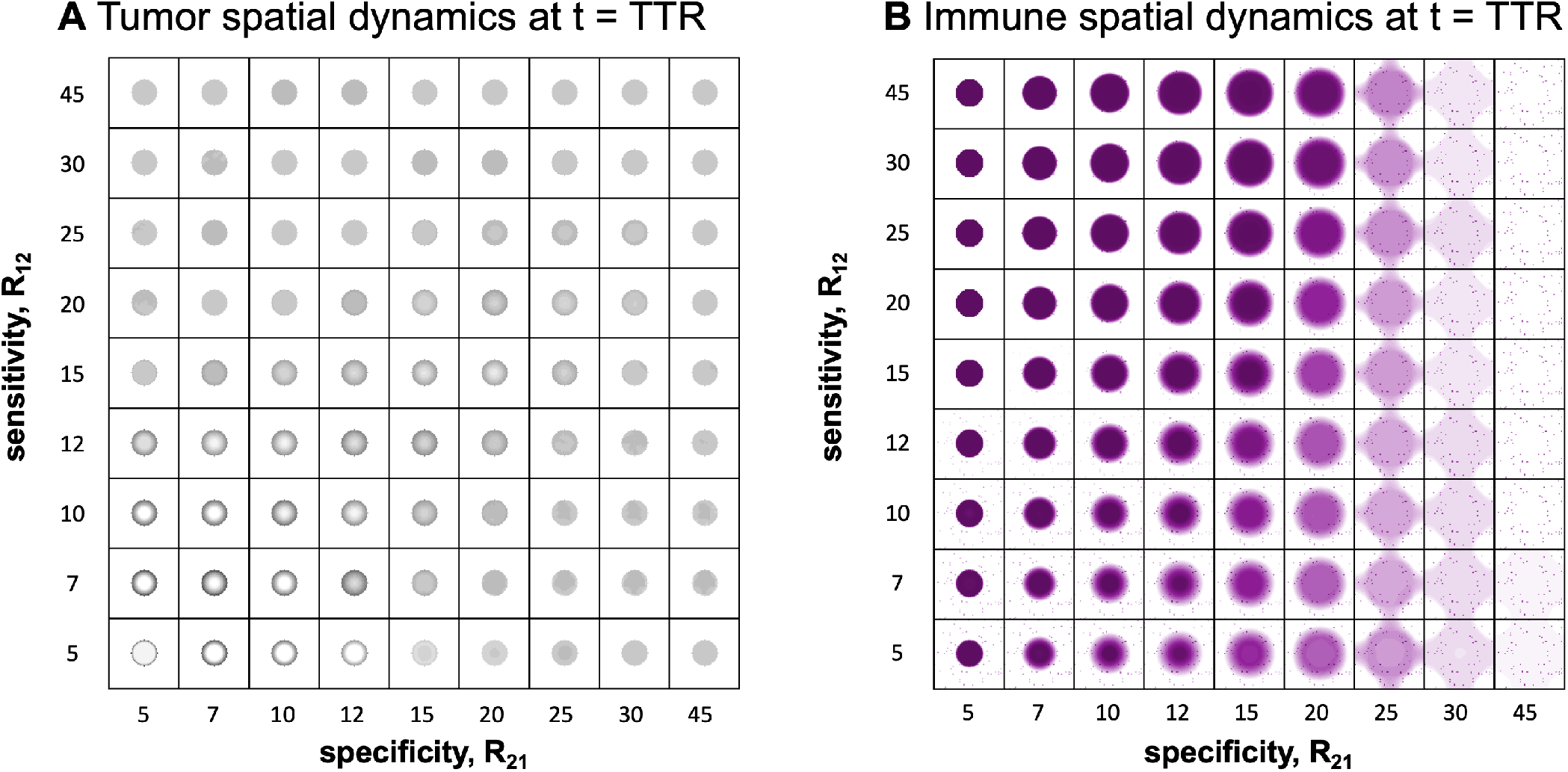
Tumor-immune spatial variegation patterns as a function of interaction kernels. Spatial maps are shown at the time to tumor regression (TTR; see figure 6) for a range of tumor-immune sensitivity (*r*_12_) and specificity (*r*_21_) values. (A) Tumor spatial maps at TTR. (B) Immune spatial maps at TTR. Spatial variegation in tumor density increases as both specificity and sensitivity are increased. As immune specificity narrows (*r*_21_ is small), immune cell concentration strongly correlates with tumor’s location with a high degree of specificity. Unless otherwise noted, parameters used are *γ* = 5, *b* = 12, *g* = 1.5, *d* = 1, *L* = 0.08 (see eqns. 6-7).

### S7.2 Stochastic Lenia

The Lenia framework can be extended to a stochastic setting by considering the likelihood of a lattice location **x** containing a single cell. Let **A**(**x**) ∈ [0, 1] be a random variable, such that the probability, *P*, of a cell potentially being added to an existing lattice location **x** at the next time step *t* + Δ*t*. This probability is determined by the Binomial distribution (if the lattice location is currently empty), at a rate of *G*(**U**(**x**))Δ*t*.

### S7.3 Particle Lenia

Particle Lenia is an extension of Lenia where movement of the particle is determined by the sum of two local gradients representing the energy field, **E**(**p**_*i*_), and migration field, **M**(**p**_*i*_). Particle *i*’s location in two-dimensional space is given by **p**_*i*_ := {*x, y*} and updated according to the following equation:

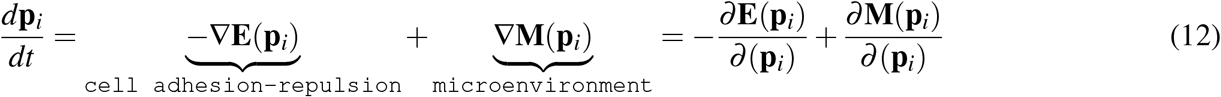

Particles (or cells) move against the local gradient of **E**(**x**), the energy field. The cell-cell repulsion and adhesion fields are a function of the potential field, **U**, such that:

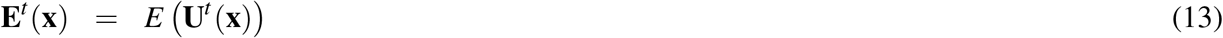

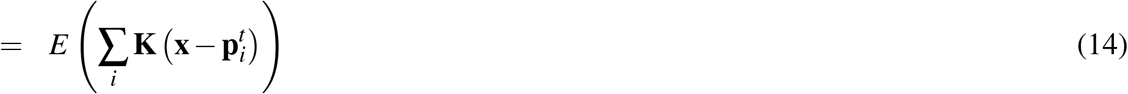

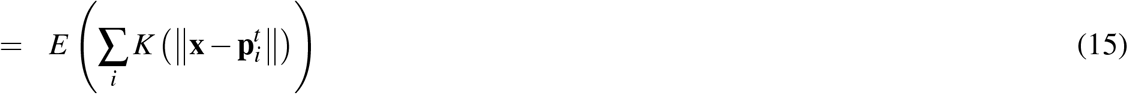

where **E**^*t*^ (**x**) is interpreted as the energy field as a function of the distance from the focal particle. Interacting cells move along the gradient of the energy field, weighted by the interaction kernel as a function of particle-particle distance, and summed over all particles.

## S8 Supplemental Video Captions

**Video S1. Supplemental Video S1: example immune infiltration simulation for perpendicular and parallel model** Video of simulations, corresponding to figure 3. Left: perpendicular model. Right: parallel model.

**Video S2. Supplemental Video S2: example immune infiltration simulation for perpendicular and parallel model** Video of simulations, corresponding to figure 3E. Simulations ordered by disease stage from left to right.

**Video S3. Supplemental Video S3: Short-range interaction kernels are more robust to Allee effects**

Spatial maps shown for deterministic Lenia (left) and stochastic Lenia (right) over time.

**Video S4. Supplemental Video S4: tumor-immune spatial variegation patterns as a function of interaction kernels** Video of simulations, corresponding to figures S1, 6E. Left: Spatial maps shown over time for a range of tumor-immune sensitivity (*r*_12_) and specificity (*r*_21_) values. Right: Immune spatial maps over time. Spatial variegation in tumor density increases as both specificity and sensitivity are increased.

As immune specificity narrows (*R*_21_ is small), immune cell concentration strongly correlates with tumor’s location with a high degree of specificity. Parameters used are *γ* = 5, *b* = 12, *g* = 1.5, *d* = 1, *L* = 0.08 (see eqns. 6-7).

## Footnotes

a For example, one might define **E** as the Lennard-Jones potential function, which describes the balance of repulsive and attractive forces as a function of the distance between two interacting particles^61^

